# Fast and Accurate EEG/MEG BEM-Based Forward Problem Solution for High-Resolution Head Models

**DOI:** 10.1101/2024.06.07.598024

**Authors:** William A. Wartman, Guillermo Nuñez Ponasso, Zhen Qi, Jens Haueisen, Burkhard Maess, Thomas R. Knösche, Konstantin Weise, Gregory M. Noetscher, Tommi Raij, Sergey N. Makaroff

## Abstract

A BEM (boundary element method) based approach is developed to accurately solve an EEG/MEG forward problem for a modern high-resolution head model in approximately 60 seconds using a common workstation. The method utilizes a charge-based BEM with fast multipole acceleration (BEM-FMM) and a “smart” mesh pre-refinement (called *b*-refinement) close to the singular source(s). No costly matrix-filling or direct solution steps typical for the standard BEM are required; the method generates on-skin voltages as well as MEG magnetic fields for high-resolution head models in approximately 60 seconds after initial model assembly. The method is verified both theoretically and experimentally.

## 1. Introduction

EEG/MEG source reconstruction, or source localization, consists of locating the neural activity within the brain from EEG/MEG-recorded measurements [1]. Many excellent open-source software packages implement EEG source localization, including Brainstorm [2], FieldTrip [3], MNE [4], and EEGLab [5]. All four packages include BEM (boundary element method)-based dipole source localization, typically using three layers extracted from the subject’s MRI: scalp, outer skull, and inner skull or “brain”. For most practical applications, the resolution of these layers is low; it is limited to less than 5-10 thousand triangles per layer. This is because a dense system matrix needs to be computed and stored in the classic BEM formulation, which becomes too large for higher-resolution accurate head models.

Recent progress in the charge-based BEM with fast multipole acceleration (a matrix-free BEM or BEM-FMM) [12] has made it possible to overcome the matrix storage limitation [13]. However, there exists one more difficulty, which is specific to the singular cortical dipole models. The default cortical surface resolution of modern head segmentation pipelines [10], [7],[11] is approximately the same as the separation distance between the dipole and the surfaces. It has long been known [14] that the BEM cannot accurately model the response of singular sources at such short distances, no matter how many neighbor surface integrals are precomputed analytically. Therefore, an adaptive mesh refinement (AMR) is necessary close to the sources [16]. While possessing an unconstrained numerical accuracy, the corresponding solution suggested in [15],[16] is still relatively slow for practical purposes.

The present study introduces a new fast method of AMR that is based on one-time “smart” mesh pre-refinement using the first, almost trivial, approximation to the solution of the BEM integral equation written in terms of the surface charge density. This method, which we name “*b*-refinement”, is quite accurate. Simultaneously, it allows us to reduce the forward solution time per dipole distribution from ca 30 min to approximately 60 seconds when a detailed head model is used. Without the costly matrix-filling and direct solution steps typical for the standard BEM, the method generates on-skin voltages as well as output MEG magnetic fields for a high-resolution (ca. 1 million facet) head model in approximately 60 seconds after its first assembly.

The study is organized as follows. Section 2 describes the idea of the method and its realization. It also describes three test cases: comparison with an analytical solution, comparison with the precise AMR algorithm [15],[16] (which is considered as the ground truth) for realistic head models with three challenging dipole locations, and finally, application to practical source reconstruction for experimental data on median nerve stimulation. Section 3 reports the obtained results in all three cases. Section 4 provides a short discussion; Section 5 concludes the study. The software is available to interested researchers via a GitHub repository [32].

## 2. Materials and Methods

### 2.1 Why adaptive mesh refinement?

Fig. 1a shows a cortical dipole placed at the posterior wall of the central sulcus halfway between the gray matter (GM) and white matter (WM) surfaces for Connectome Young Adult subject 110411 [6]. The dashed line indicates the dipole’s orientation. The head model was segmented by headreco [7] and has approximately 1 million triangular facets with an average edge length of approximately 1.5 mm (GM) and 1.4 mm (WM).

**Fig. 1.**
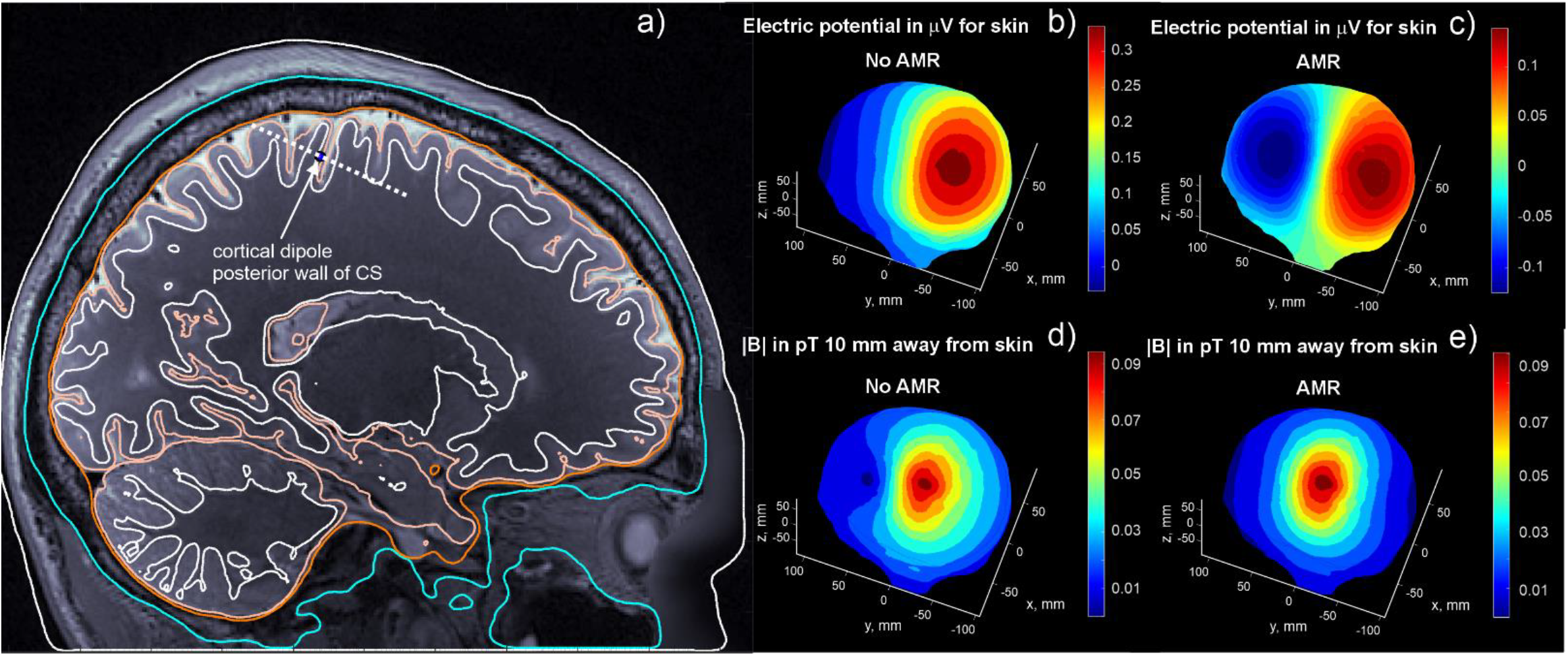
a) Cortical dipole position at the posterior wall of the central sulcus. b,c) On-skin electric potential with and without adaptive mesh refinement. d, e) Magnitude of the magnetic field 10 mm away from the skin surface with and without adaptive mesh refinement. The cortical dipole has a moment of 4e-9 A·m.

Such an edge length (triangle size) – the default computational resolution of the common segmentation packages FreeSurfer [10], headreco [7], and Charm [11] – is typically greater than or equal to half of the cortical thickness [8],[9], and therefore also greater than the separation of the mid-surface dipole from either cortical boundary. This resolution is insufficient to enable a BEM-based solution to accurately model the singular dipole source very close to the nearest “large” triangles. As a result, inaccurate predictions are obtained for both the on-skin electric potential (Fig. 1b) and off-skin magnetic field (Fig. 1d) as compared to those for a sufficiently refined surface mesh in Fig. 1 c, e, respectively. There, the refined facets in the vicinity of the source are approximately 10-20 times smaller than the separation distance between GM and WM shells (cortical thickness).

Therefore, regardless of the initial segmentation, mesh refinement must be performed for proper EEG/MEG source modeling. Due to present computer hardware limitations, it is not practical to employ a uniform (global) mesh refinement scheme that achieves the requisite resolution. A local, and ideally adaptive (based on physically justified criteria) mesh refinement is therefore required for accurate EEG/MEG source modeling by BEM.

### 2.2 Existing h-refinement method

The charge-based BEM equation for the surface charge density *ρ*(***r***) induced at all interfaces *S* of a piecewise-homogeneous multi-compartment head model is written in the following form [12]

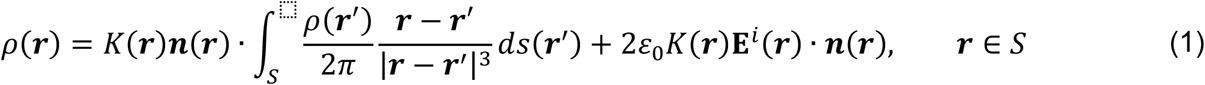

where 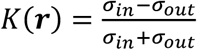 is the electric conductivity contrast for the facet positioned at ***r, n***(***r***) is the outer normal vector at the compartmental interfaces, and *σ*_*in*_ and *σ*_*out*_ are the conductivities of the materials just inside and outside (respectively) the interface at ***r***. The induced charges are generated by the impressed or primary electric field **E**^*i*^(***r***) of a cortical current dipole (or a cluster thereof). A solution *ρ*(***r***) to Eq. (1) is found iteratively via the Generalized Minimum Residual Method (GMRES), which takes its initial estimate for *ρ*(***r***) as:

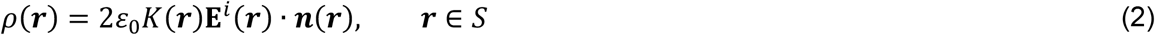

In a previous approach [15],[16], an *h*-refinement method was applied wherein facets of the model were selected for 4:1 barycentric subdivision according to the absolute value of total charge *q*_*m*_ = |*A*_*m*_*ρ*_*m*_| upon them, where *A*_*m*_ and *ρ*_*m*_ denote the area and charge density, respectively, on the facet with index *m*. The solution was carried out in alternating steps of “solve” (by a full GMRES solution) and “refine” until a convergence criterion for relative inter-iteration change in skin potential was reached. While the method produced quite accurate results [16], the runtime became prohibitive when looping over multiple configurations of sources.

### 2.3 Concept of b-refinement

In this study, we suggest using the initial estimate for *ρ*(***r***) by Eq. (2) for an *a priori* mesh refinement, which is performed only once and before carrying out the iterative GMRES solution. While this initial estimate does not take into account the final charge redistribution due to self-interaction, it sufficiently identifies regions where *q*_*m*_ will have the maximum absolute values. These regions are indeed located close to the source(s), but their exact topology depends on the compartmental conductivities and interfacial bends. This method is used in place of *h*-refinement, although a multi-phase process that combines the two may be viable.

We refer to the mesh refinement method based on Eq. (2) as “*b*-refinement” to highlight its association with the iterative solution of a well-conditioned system of linear equations in the form *x* + *Âx* = *b* where the first approximation to *x* is *x* = *b* (the right-hand side). This is the form taken by Eq. (1) when discretized; for a detailed explanation, see [16].

### 2.4 Mesh refinement strategy

First, Eq. (2) is used to find the approximate surface charge distribution on the conductivity interfaces. Following Refs. [15],[16], the facets having the largest *total* charges *q* are each subdivided into four congruent triangles whose edges are the halves of the original edges. To restore manifoldness after refinement, the border facets of the refinement region are also subdivided into two facets each.

An *m*-th facet is refined if its total charge *q*_*m*_ satisfies the following inequality:

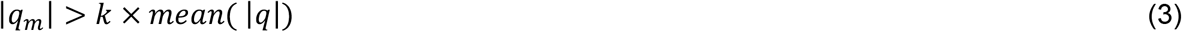

where *k* is a constant on the order of 5-10 and the absolute mean charge value is found over all facets. The refinement is performed iteratively according to the following steps: find a new initial charge distribution estimate following Eq. (2), select facets for refinement according to Eq. (3), subdivide those facets, and repeat. The number of necessary mesh refinement steps was empirically found to vary from 3 (the sphere model) to 4-5 (realistic head models). Surface-preserving Taubin smoothing [17] is additionally applied at every refinement step, with the scale factors of -0.62 and +0.60, respectively. After application of *b*-refinement, the full charge solution is found by GMRES as usual, without further mesh refinement steps.

### 2.5 Validation example - a four-layer sphere

A classical EEG and MEG solution for a four-layer conducting sphere shown in Fig. 2 is analyzed first, similar to many other studies (cf. [18],[19]).To validate and compare the accuracy of the *b*-refinement approach, we create the standard four-layer sphere model, where analytical solutions for infinitely short EEG [20],[21] and MEG [22] dipoles exist and can be used as a reference. We test both horizontal (challenging for EEG) and vertical (challenging for MEG) dipoles located 2 mm away from the “brain” surface in Fig. 2. To test different mesh resolutions, we create and clone nine high-quality triangular sphere meshes with numbers of triangular facets ranging from approximately 6,000 to 0.4 M using a high-quality surface mesh generator developed in [23],[24].

**Fig. 2.**
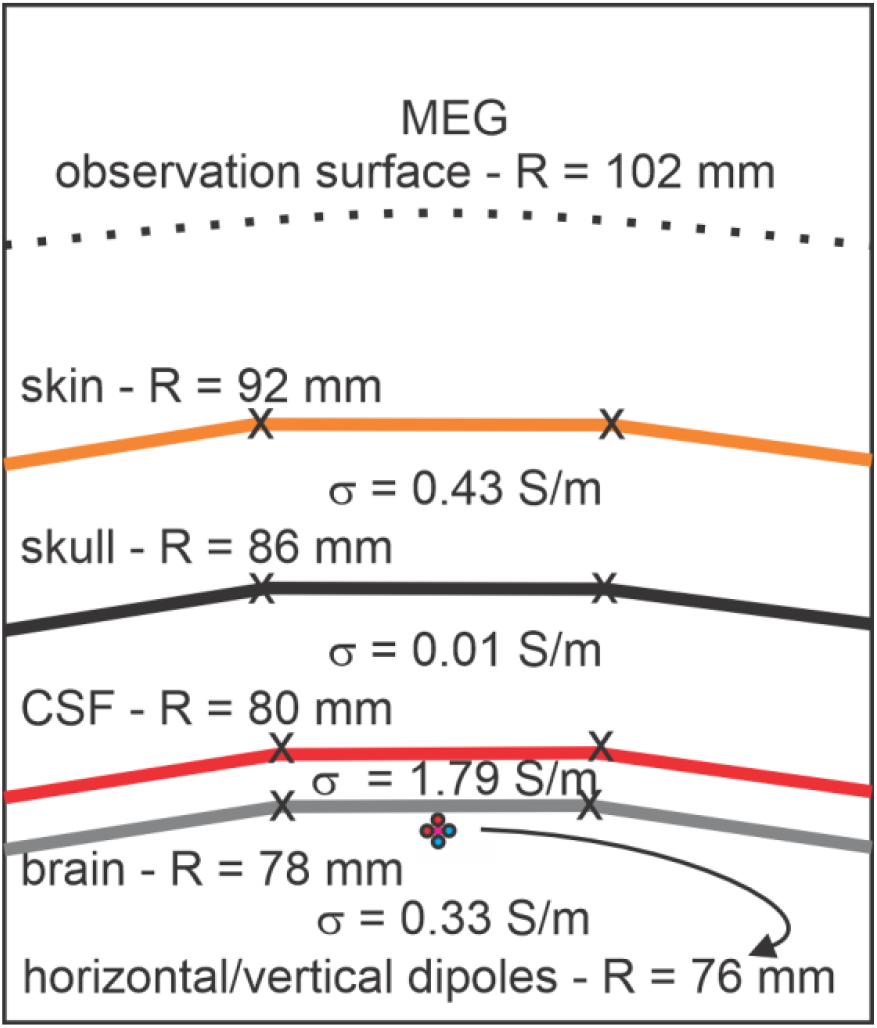
Model of four-layer sphere used for comparison along with the dipole positions and conductivity values.

### 2.6 Validation example - dipole sources in realistic head models

Headreco models for Connectome Young Adult [6] subjects 110411 and 120111 are modeled with three dipole locations each: posterior wall of central sulcus, *M*_1HAND_ area, and auditory cortex. The dipole locations for subject 110411 are shown in Fig. 3. The material properties are those employed by the open-source software SimNIBS [7].

**Fig. 3.**
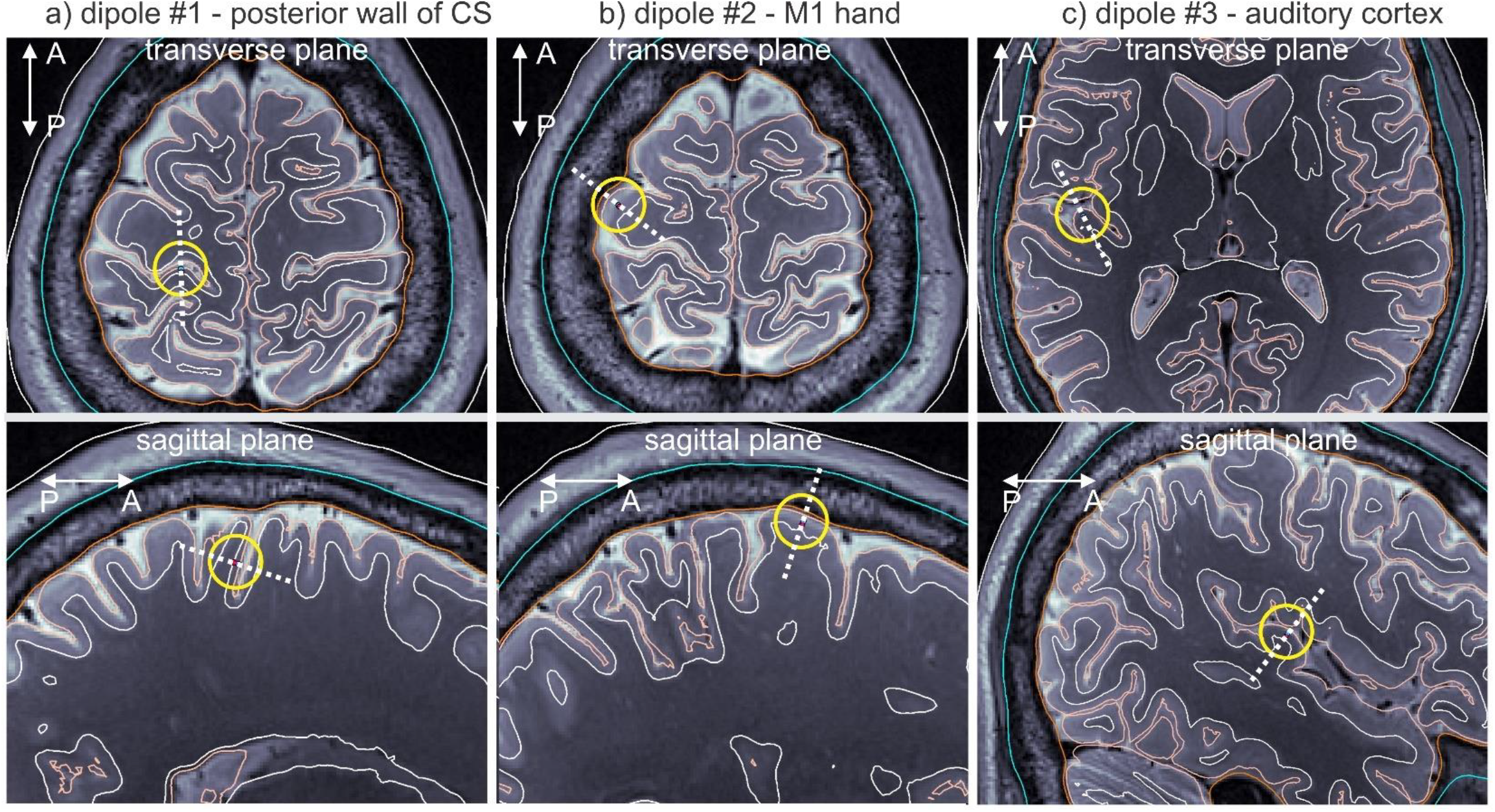
Three dipole positions used for comparison purposes. a) Position at the posterior wall of the central sulcus. b) Position within *M*_1HAND_ area. c) Position within auditory cortex.

Both EEG (on-skin voltage) and MEG (vector magnetic field 10 mm away from skin) outcomes are compared across two solutions: the *b*-refinement method and the most accurate adaptive mesh refinement solution [15],[16], which shows excellent self-convergence but runs approximately 30 times slower. The standard 2-norm error and the RDM (relative difference measure) error (cf. [16]) are employed for comparison.

### 2.7 Source localization with b-refinement method from experimental data

A 1 mm T1 MRI scan of a healthy subject at A.A. Martinos Ctr. for Biomedical Imaging, Massachusetts General Hospital was followed by median nerve stimulation. Electrical stimuli over the median nerve at the right wrist were delivered to the subject using brief transcutaneous pulses every 1.5 seconds. The task was to respond to each stimulus by pushing a button with the left-hand index finger. This generates EEG evoked responses in *S*_1HAND_ (primary somatosensory cortex contralateral to the nerve stimuli), and *M*_1HAND_ at different latencies [25],[26].

The P20/N20 response peaking at about 20 ms after the stimulus in Fig. 4a was used for source localization, since at this latency its neuronal generators are well-known to be located in the posterior wall of the central sulcus in the Brodmann area 3b [27],[28]. The recordings (3 runs of 80 trials each) were done at Massachusetts General Hospital using a 70 channel EEG system with electrode locations shown in Fig. 4b and the P20/N20 normalized electrode voltages shown in Fig. 4c. The head model and cortical surface reconstruction were obtained with SimNIBS headreco segmentation.

**Fig. 4.**
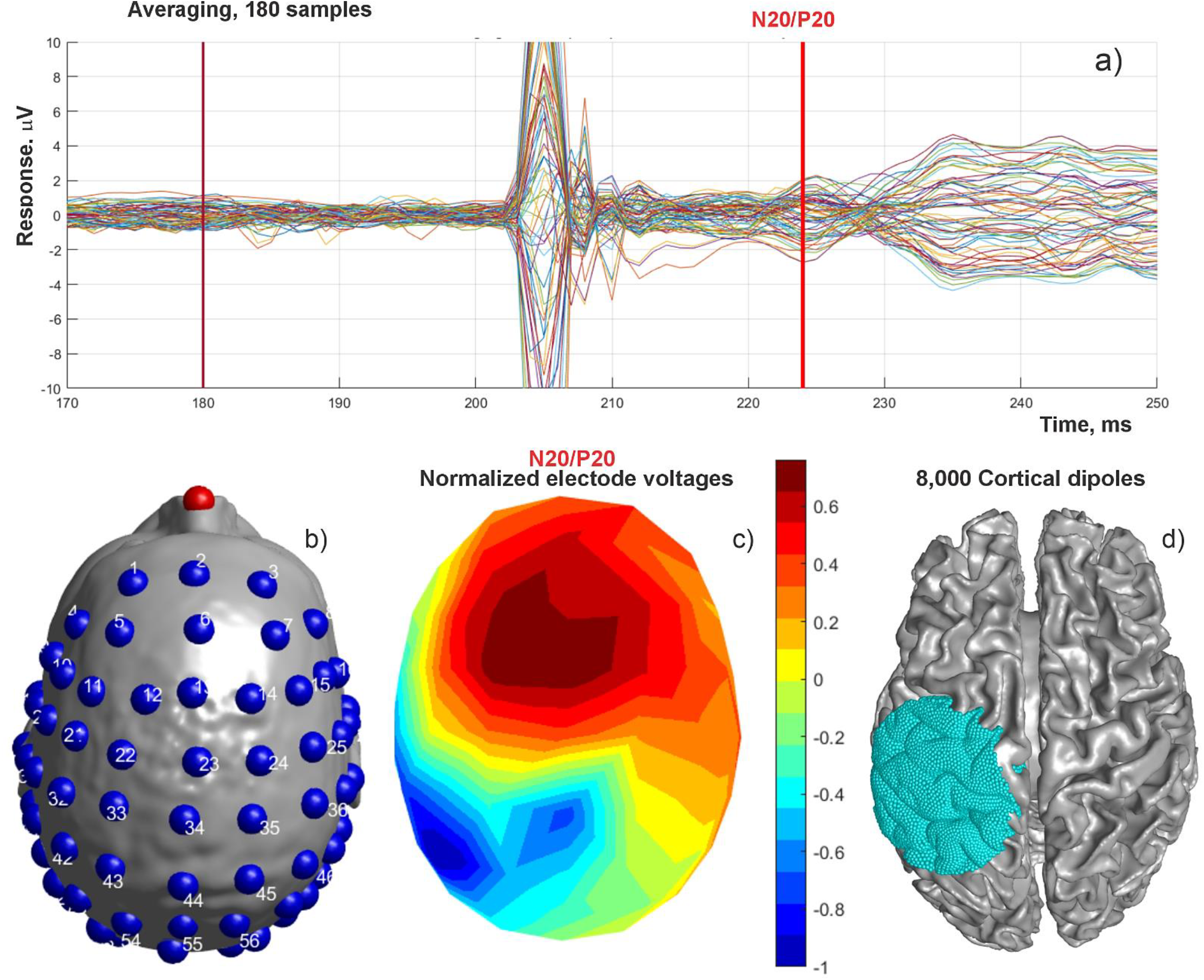
a) Electrode voltages and P20/N20 peak after median nerve stimulation of a heathy subject. b) Electrode positions (70 electrodes). c) Normalized on-skin voltage distribution for P20/N20 peak. d) Positions of 8,000 cortical dipoles used for source localization.

The *b*-refinement method was used to find forward solutions for 8,000 cortical dipoles located in the vicinity of the expected neuronal response, at the mid-surface between gray and white matter. The dipole locations are schematically shown in Fig. 4d. The source localization problem was solved both by the standard Moore-Penrose pseudoinverse and by computing a minimum norm least-squares solution; both methods are implemented in MATLAB. The electrode voltages are normalized by their self-variances. The goal of this experiment is to compare the known anatomical region of the neural activity with the modeling predictions.

## 3. Results

### 3.1 Validation example - a four-layer sphere

Fig. 5 demonstrates the solution process for a coarse four-layer sphere model. The source in this case was the horizontal electric dipole in Fig. 2 with a moment of 4e-11 A/m. The figure’s first column shows, for each shell, the initial surface charge distribution estimates from Eq. (2) that were used as the starting point for AMR. The second column demonstrates the corresponding meshes obtained after four *b*-refinement steps, also for every shell. Note that the fourth step produces triangles that are barely visible on the figure. The figure’s third column demonstrates the resulting forward EEG solution (potential on every shell) after AMR is done.

**Fig. 5.**
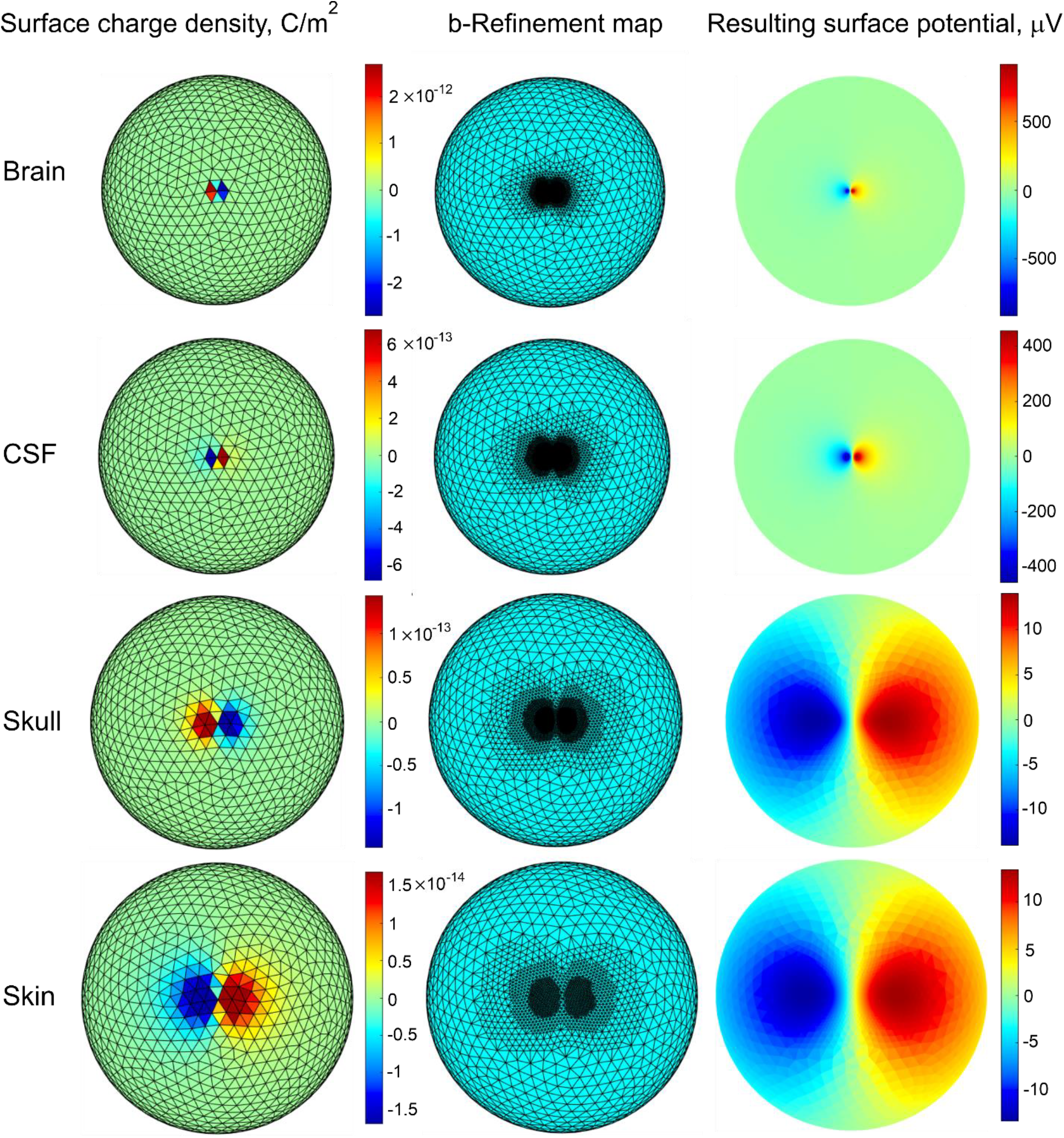
First column – initial surface charge distributions for a coarse four-layer sphere model from Fig. 2. Second column – the corresponding b-refinement maps obtained with the four refinement steps for every shell. Third column – surface electric potential for every shell obtained after mesh refinement. All data are for the horizontal electric dipole in Fig. 2 with the moment of 4e-11 A·m.

The obtained solution was compared with the analytical solution as described in the previous section. Fig. 6a,b shows the 2-norm/RDM error for electric potential over the entire outer shell (skin surface), and Fig. 6c,d shows the 2-norm/RDM error in the vector magnetic field at a shell 10 mm from the skin surface. The results shown in Fig. 6 a,b,c,d are for the horizontal electric dipole 2 mm away from the innermost shell (the “brain” surface), while Fig. 6 e,f,g,h presents results for the vertical electric dipole in the same location (cf. Fig. 2).

**Fig. 6.**
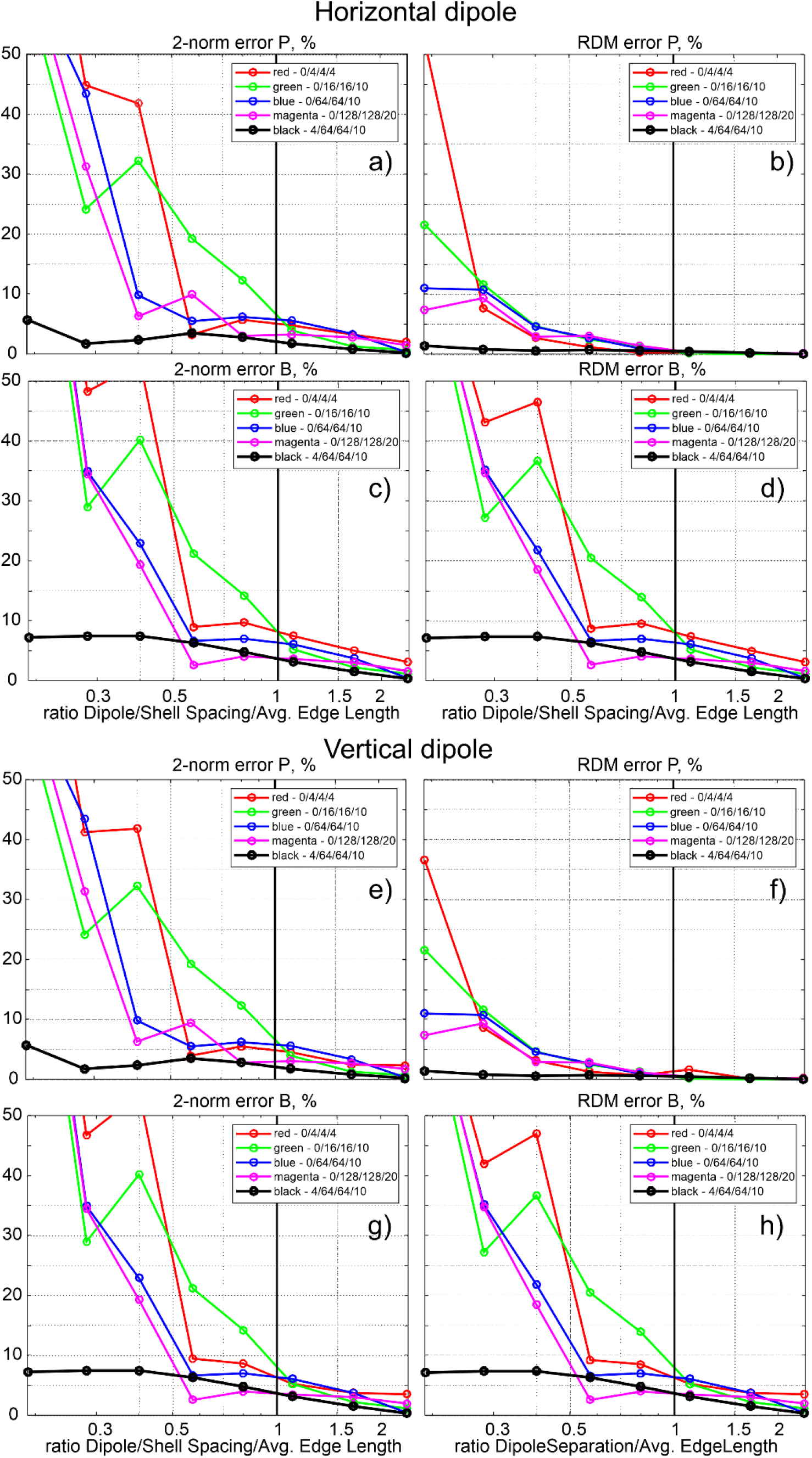
2-norm and RDM error percentages between analytical and numerical solutions for the four-layer sphere model. Results for the b-refinement with four steps (levels) are black curves.

Every plot in Fig. 6 has five curves. The red, green, blue, and magenta curves are those without adaptive mesh refinement. The red curve is the solution with analytical integration over 4 neighbor facets for both the field and the potential [12]; the green curve is the solution with analytical integration over 16 neighbor facets; the blue curve utilizes 64 neighbor-facet integrals; and the magenta curve uses 128. The black curve is the *b*-refinement solution with four refinement steps as shown in Fig. 5. This solution utilizes 64 neighbor integrals.

In every pane of Fig. 6, the argument is the dimensionless ratio of the dipole distance from the “brain” surface (2 mm) or the spacing between the “CSF” shell and the “brain” shell (also 2 mm) to the average edge length of the non-refined, original model mesh. The last number in every curve legend is a dimensionless radius of a sphere (in terms of average edge length) within which an analytical integration of the dipole’s primary electric field [30] in Eq. (2);

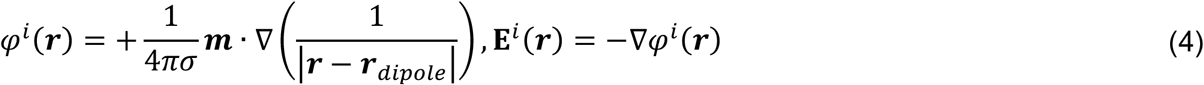

over the planar surface triangles is performed instead of the center-point approximation used anywhere else. Here, ***m*** = *I*_0_***d*** is the vector current dipole moment (A·m), ***d*** is the vector from the current sink −*I*_0_ to the current source +*I*_0_, and *σ* is the background conductivity of the medium where the dipole source is located.

### 3.2 Validation example - dipole sources in realistic head models

Fig. 7 demonstrates the *b*-refinement method outcome for the realistic head topology. Fig. 7a is the original headreco segmentation superimposed onto T1 NIfTI data for subject 110411. The dipole position at the posterior wall of the central sulcus is marked by a circle. Red dots indicate edge intersections with the transverse plane. Fig. 7b shows the mesh for the gray matter surface close to the dipole position after 4 *b-*refinement steps. Refinement level 4 is deeply inside the sulcus and is not visible. Fig. 7c is the same plot as in Fig. 7a, but after *b*-refinement with four steps. Fig. 7d shows the mesh for the white matter surface close to the dipole position after 4 *b-* refinement steps. The skin surface is not refined for the present dipole position when four refinement steps are used. The *b*-refinement leads to a moderate total mesh size increase of approximately 13% (1.04 M to 1.17 M facets) in the present case.

**Fig. 7.**
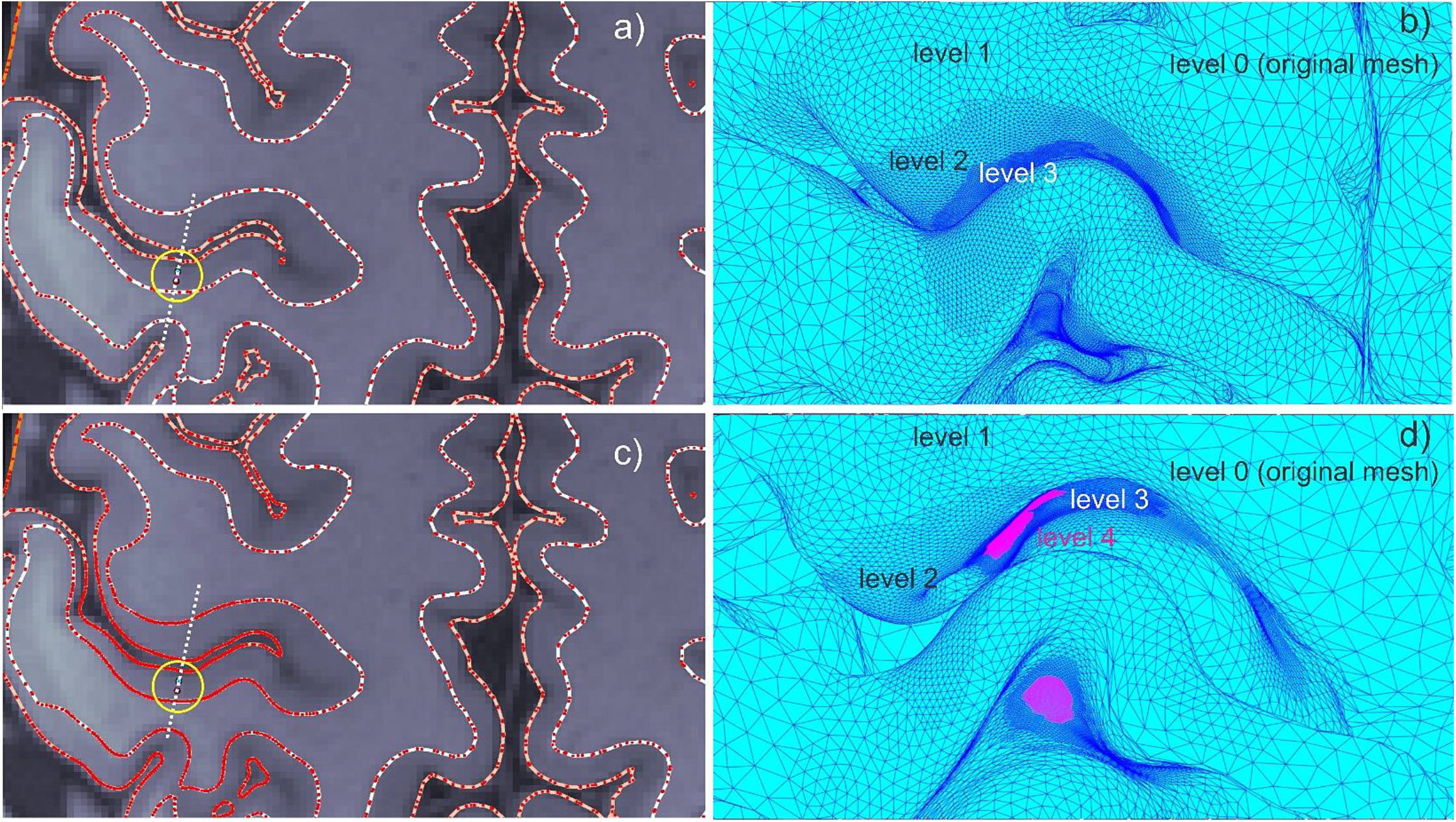
a) Original headreco segmentation superimposed onto T1 NIfTI data for subject 110411. The dipole position at the posterior wall of the central sulcus is marked by a circle. Red dots indicate edge intersections with the transverse plane. b) *b*-refinement for the gray matter surface close to the dipole position after 4 refinement steps. Refinement level 4 is deeply inside the sulcus and is not visible. c) The same plot as in a), but after *b*-refinement with four steps. d) *b-*refinement for the white matter surface close to the dipole position after 4 refinement steps.

Table 1 reports averaged 2-norm and RDM error percentages for the two head models and three dipole positions (previous section, Fig. 3) between the *b*-refinement method with four levels of refinement and the accurate converging AMR solution [16],[31]. For the EEG electric potential, the errors are computed for the entire skin surface. For the MEG vector magnetic field (magnetic flux) **B**, the errors are computed 10 mm away from the skin surface in the normal direction.

**Table 1.**
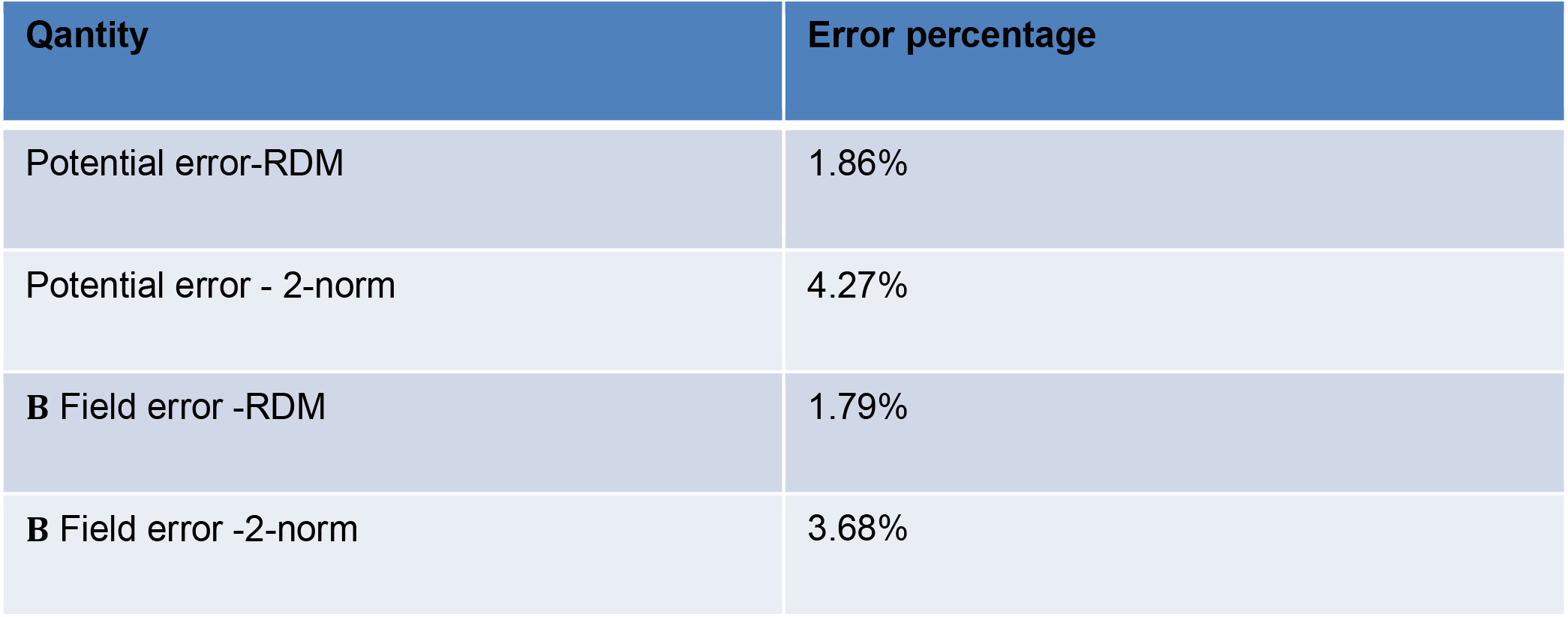
Averaged 2-norm and RDM error percentages for two head models and three dipole positions (Fig. 3) between the *b*-refinement method with four levels of refinement and the accurate self-converging AMR solution [16],[31]. For the EEG electric potential, the errors are computed for the entire skin surface. For the MEG vector magnetic field (magnetic flux) **B**, the errors are computed at a surface 10 mm away from the skin surface (in the normal direction).

Fig. 8 demonstrates the dipole fields within cortical as well as extracerebral compartments for subject 120111. The first row in Fig. 8 is the electric potential distribution in three principal planes for subject 120111. The cortical dipole is located at the posterior wall of the central sulcus. The second row in Fig. 8 is the magnetic field (flux) magnitude distribution for the same cortical dipole in three principal planes. Note that a logarithmic scale has been used in both cases.

**Fig. 8.**
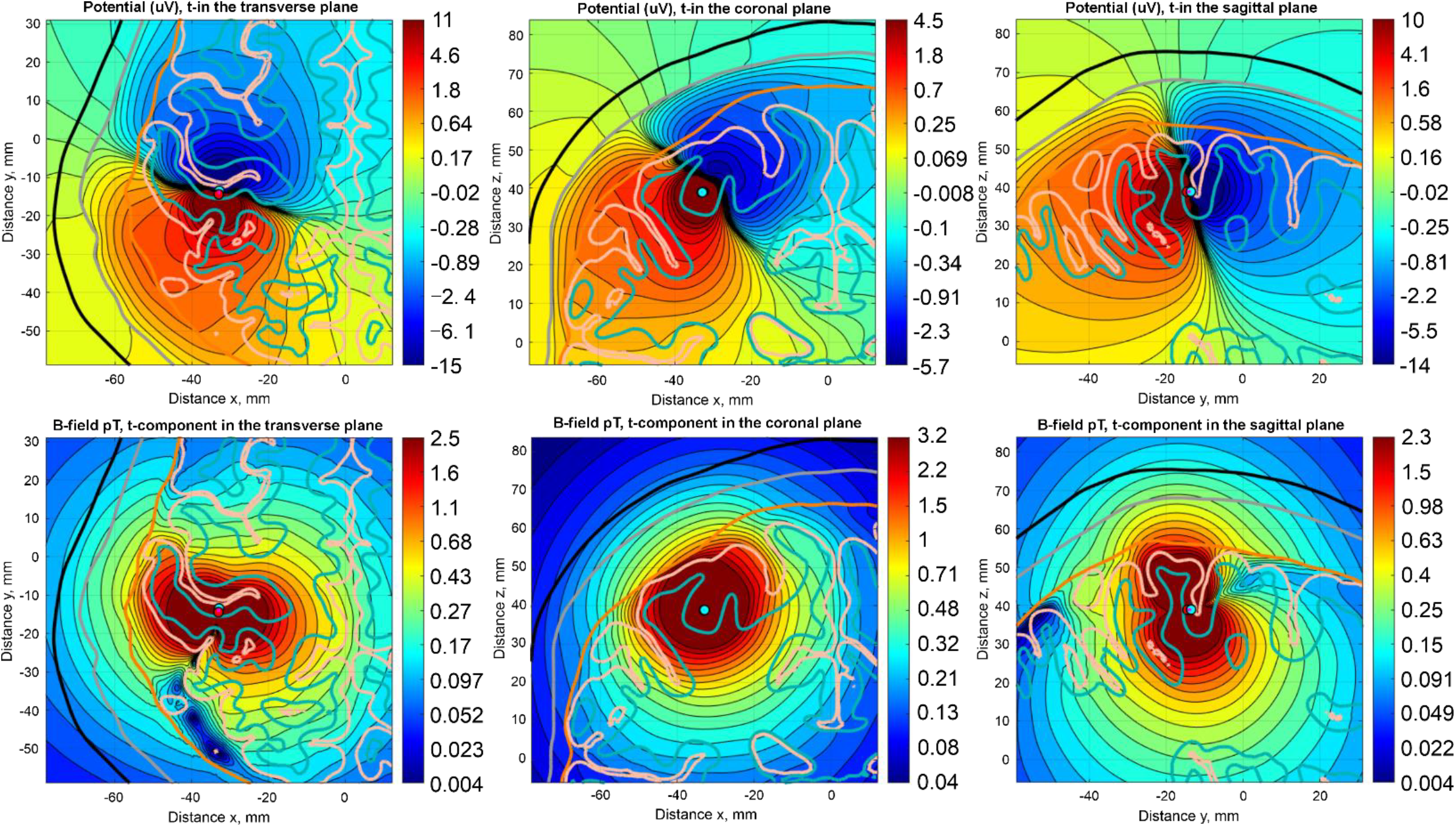
First row – electric potential distribution in three principal planes for subject 120111. The cortical dipole is located at the posterior wall of the central sulcus. Second row – magnetic field (flux) magnitude distribution for the same cortical dipole in three principal planes. Note that a logarithmic scale is used in both cases.

### 3.3 Source localization with b-refinement method using experimental EEG data

Basic source reconstruction was performed for the experimental data on median nerve stimulation described in Section 2.7. Fig. 9 displays the reconstructed dipole strength density for the P20/N20 peak. Red spheres indicate cortical dipole locations with the maximum positive strength (dipoles directed from white matter to gray matter) while blue dots indicate cortical dipoles with the maximum negative strength (dipoles directed oppositely). The three panes of Fig. 9 correspond to the threshold values of the dipole strength set to 70^th^, 80^th^, and 90^th^ percentiles. The crown of the postcentral gyrus is indicated by a black curve in every pane.

**Fig. 9.**
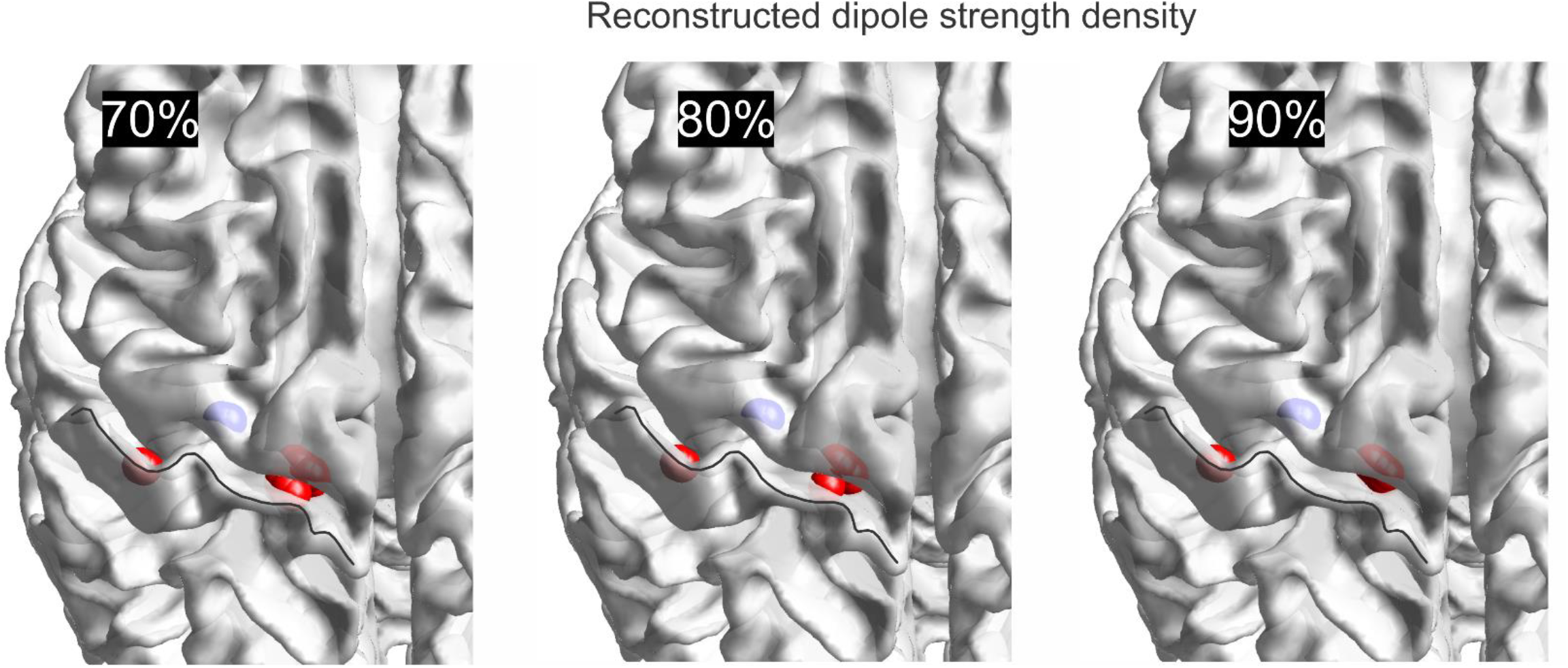
Reconstructed dipole strength density for the experimental data on median nerve stimulation. Red spheres indicate cortical dipoles with the maximum positive strength (directed from white matter to gray matter) while blue dots indicate cortical dipoles with the maximum negative strength (directed oppositely). From left to right: threshold values of dipole strength are set up at 70^th^, 80^th^, and 90^th^ percentiles. Crown of the postcentral gyrus is indicated by a black curve in every pane.

The results obtained with the Moore-Penrose pseudoinverse and the minimum-norm least-squares solution were found to be nearly identical.

## 4. Discussion

### 4.1 Validation example - a four-layer sphere

The results given in Fig. 6 demonstrate that the *b*-refinement method leads to much lower and very consistent computational errors for relatively coarse meshes, when the ratio of the dipole distance from the nearest shell to the average edge length of the non-refined (original) model mesh is less than one. Exactly this case is observed for the realistic head models. A larger number of neighbor integrals computed analytically also reduces the error, but this approach becomes impractical due to the large storage and extensive precomputations necessary.

In a further series of numerical experiments, “segmentation noise” was also introduced by randomly perturbing mesh vertices, and excellent resilience against the noise was observed when using *b*-refinement.

### 4.2 Validation example - dipole sources in realistic head models

Table 1 indicates that the RDM error for both EEG and MEG forward problems does not exceed 2% on average when the *b*-refinement is applied with a relatively small number of steps. Similarly, the average 2-norm error does not exceed 4.5%. This is certainly acceptable in practical applications. Notably, the *b*-refinement results in a very modest overall mesh size increase of 13-15% or so for a point dipole or a small cluster of closely spaced dipoles up to 5-7 mm in size. Further, Fig. 8 demonstrates that the dipole field could be substantially distorted by nearby white and gray matter interfaces. This underscores the importance of the high-resolution models and the adaptive mesh refinement for source reconstruction.

### 4.3 Source localization with b-refinement method using experimental EEG data

The source reconstruction maps given in Fig. 9 indicate that the present approach provides localization results for median nerve stimulation that agree with the experimental predictions for neuronal generators [27],[28]. All three panes in Fig. 9 predict source locations in the posterior wall of the central sulcus or at its bottom, in the Brodmann area 3b. Those are the red spheres in Fig. 9.

At the same time, dipolar sources of opposite polarity can also be predicted at the anterior wall of the central sulcus as shown by a blue cluster in Fig. 9. This result is to be expected since the simultaneous interchange of both the wall and the polarity would lead to nearly the same dipole field. A (small) change in the source location has apparently little effect on the ill-posed EEG inverse problem. In any case, the solution remains stable with regard to the source strength threshold – all three panes in Fig. 9 are quite similar to each other.

## 5. Conclusion

The *b*-refinement method for forward EEG and MEG problems introduced in this study has been verified both theoretically and experimentally. This method, in conjunction with the boundary element fast multipole method (BEM-FMM), allows us to solve a forward problem for a single dipole or a compact dipole cluster in approximately 60 seconds when a modern detailed head model is used. Without the costly matrix-filling and direct solution steps typical for the standard BEM, the method generates on-skin voltages as well as output MEG magnetic fields for a high-resolution (ca. 1M facet) head model in approximately 60 seconds after its first assembly.

## Acknowledgements

WAW, GNP, ZQ, GMN, and SNM were supported by the NIBIB grant R01EB035484 and NIMH grant R01MH130490. TR was partially supported by the NINDS grant 1R01NS126337. KW was partially supported by the BMBF grant: 01GQ2201. JH received funding from the German Federal Ministry of Education and Research (BMBF) grant Dry-Pole (01GQ2304A) and the Free State of Thuringia (2018 IZN 004), co-financed by the European Union under the European Regional Development Fund (ERDF).

